# Identification of Candidate Drugs for Heart Failure using Tensor Decomposition-Based Unsupervised Feature Extraction Applied to Integrated Analysis of Gene Expression between Heart Failure and DrugMatrix Datasets

**DOI:** 10.1101/117465

**Authors:** Y-h. Taguchi

**Affiliations:** Department of Physics, Chuo University, Tokyo 112-8551, Japan

**Keywords:** Tensor decomposition, Drug discovery, Heart diseases

## Abstract

Identifying drug target genes in gene expression profiles is not straightforward. Because a drug targets not mRNAs but proteins, mRNA expression of drug target genes is not always altered. In addition, the interaction between a drug and protein can be context dependent; this means that simple drug incubation experiments on cell lines do not always reflect the real situation during active disease. In this paper, I apply tensor decomposition-based unsupervised feature extraction to the integrated analysis of gene expression between heart failure and the DrugMatrix dataset where comprehensive data on gene expression during various drug treatments of rats were reported. I found that this strategy, in a fully unsupervised manner, enables us to identify a combined set of genes and compounds, for which various associations with heart failure were reported.

## 1 Introduction

*In silico* drug discovery is an important task because experimental identification/verification of therapeutic compounds is a time-consuming and expensive process. There are two major trends of *in silico* drug discovery: the ligand-based approach [1] and structure-based [2] approach. The former is very straightforward; new drug candidates are identified based upon the similarity with known drugs no matter how the similarity is evaluated. Although it is a powerful method, there are some drawbacks; if there are no known drugs for target proteins, then there is no way to find new drug candidates. Even if there are many known drugs for the target protein, it is hopeless to find compounds that are effective but without any similarity with known drugs. The second, structure-based approach, can address these weaknesses. It can identify new therapeutic compounds even without the information about known drugs. Of course, there are some drawbacks in this strategy, too. If the target protein structure is not known, it must be predicted prior to the drug discovery process. Even if the target protein’s structure is known, because we need to numerically verify the binding affinity between the ligand compound and target protein, which also requires a large amount of computational resources, structure-based *in silico* drug discovery is still far from easy to perform. In addition, prediction accuracy of protein structure and of ligand-binding structure is not very high at all. Thus, it would be helpful to have an additional/alternative strategy for *in silico* drug discovery.

Recently, an alternative approach was proposed that is aimed at finding drug candidates computationally using gene expression profiles of cell lines treated with compounds [3, 4]. This third approach is not straightforward at all. First of all, because compounds target not mRNAs but proteins, mRNA expression of drug target proteins is not always affected. Therefore, direct identification of a drug target protein in gene expression data cannot be done. Second, gene expression alteration caused by treatment with a compound may be context dependent; in other words, in a cell line, the gene expression difference caused by incubation with a compound may differ from that in diseases. To compensate these difficulties, the gene expression signature strategy was developed. In this approach, gene expression alteration profiles caused by treatment of a cell line with various drug candidates are compared with those of known drugs. If the profiles are similar, then new drug candidates are expected to function similarly to known drugs. Although this third strategy is a useful one, if there are no known drugs for the target disease, this approach cannot function at all as in the case of ligand-based approaches.

Some examples aiming to identify drug-target interaction from gene expression based upon previous knowledges are as follows. Wang et al [5] tried to identify on and off-targets of drugs based upon similarity between drug-induced *in vitro* genomic expression changes. Iwata et al [6] explored potential target proteins with cell-specific transcriptional similarity using chemical–protein interactome. Lee et al [7] tried drug repositioning for cancer therapy based on large-scale drug-induced transcriptional signatures. Although these are only a few examples, they need pre-knowledge about drugtarget interactions. Alternatively, instead of drug-target interactions, drug-disease interaction is investigated. For example, Cheng et al [8] tried to measure the connectivity between disease gene expression signatures and compound-induced gene expression profiles. Sirota et al [9] also integrated gene expression measurements from 100 diseases and gene expression measurements on 164 drug compounds, yielding predicted therapeutic potentials for these drugs. Iorio et al [10] investigated compound-targeted biological pathways based upon gene expression similarity. They are unsupervised approaches to some extent, but target genes cannot be exploited.

In this paper, I propose a strategy that can infer drug candidates from drug treatment-associated gene expression profiles without the information about known compounds for diseases. In this strategy, I employ the tensor decomposition (TD)-based unsupervised feature extraction (FE) approach, which is an extension of the recently proposed principal component analysis (PCA)-based unsupervised FE; PCA-based unsupervised FE successfully solved various bioinformatic problems [11–29]. In this TD-based strategy, tensors were generated using a mathematical product of a gene expression profile of drug-treated cell lines and of a gene expression profile of a disease. Then, pairs of compounds and genes are identified whose mRNA expression alteration is associated with drug-treated cell lines and is coincident with such alteration during disease progression. Biological evaluation of the identified genes and compounds based upon past studies turned out to be promising.

## 2 Materials and Methods

### 2.1 Mathematics of TD

In this subsection, I briefly discuss what the TD is and how I apply TD to the present problem. Suppose an *m*-mode tensor 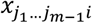 represents gene expression of the *i*th gene under the *j*_k_ (*k*=1,…,*m*-1, *j*_k_=1,…,*N*_k_) conditions, examples of which are diseases, patients, tissues, and time points. Then, TD is defined as

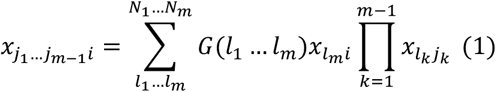

where *G*(*l*_1_ … *l_m_*) is a core tensor and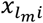 and 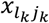 are singular value matrices that are supposed to be orthogonal to one another. Because *G*(*l*_1_ … *l_m_*) is assumed to be as large as 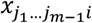, it is obviously an overcomplete problem; thus, there are no unique solutions. To solve TD uniquely, I specifically employed the higher-order singular value decomposition [30] (HOSVD) algorithm that tries to attain TD such that smaller number of core tensors and singular value vectors can represent 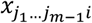 as much as possible.

### 2.2 Tensor Generation for Integrated Analysis

It is quite common when there is a set of gene expression profiles of human cell lines or model animals treated with various compounds at multiple dose densities. For example, Drug Matrix^1^ and LINCS [31] are good examples, although the former comprises only temporal gene expression after drug treatments. Nonetheless, it is not easy to infer a drug’s action on diseases by means of only these gene expression profiles; some kind of integrated analysis with disease gene expression profiles is required, but it is not so straightforward. Candidate drugs should satisfy these conditions:

- Gene expression in these profiles must significantly decrease or increase with the increasing dose density of compounds.
- Gene expression alteration caused by drug treatment must be significantly coincident with that associated with disease progression.

How these two independent significance values can be evaluated is unclear. For example, we can have two sets of significant gene expression alterations of the *i*th gene, {Δ*x_i_*}, caused by drug treatment and those of the *i*’th gene, {Δ*x*′_*i*′_}, during disease progression, respectively. First, we need to test whether the two sets of genes are significantly overlapping. Next, when there is a significant overlap, we have to determine whether these two gene expression alteration profiles correlate significantly. Furthermore, because the analysis is usually conducted among multiple compounds, all the significance evaluation must be corrected based upon a multiple comparison criterion. It is obviously a complicated and not a promising strategy.

Nevertheless, if we can have gene expression profiles expressed via a tensor, 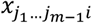, where *j*_k_ (*k*=1,…,*m*-1) corresponds to drug candidates, dose density, and disease progression, we can easily evaluate a candidate drug using TD, eq. (1). If there are 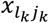 values that represent significant dependence upon dose densities and disease progression, genes’ and compounds’ singular value vectors that share core tensor *G* with larger absolute values with these 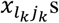 can be used for the selection of genes as well as compounds as follows.

Suppose {*l*_*k*_ } is a set of indices of genes’ or compounds’ singular value vectors that are associated with significant dose density dependence as well as disease progression dependence. Genes and compounds can be identified as being associated with significant singular value vector components. For this purpose, *P*-values are attributed to each *i*th gene/*j*_*k*_ th compound assuming a *χ*^2^ distribution,

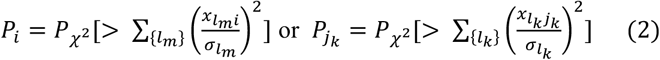

Where 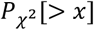 is the cumulative probability that the argument is greater than *x* assuming the *χ*^2^ distribution and 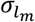 and 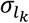 are standard deviation. After adjusting *P*-values using the Benjamini–Hochberg (BH) criterion [32], genes and compounds that have significant P-values, e.g., less than 0.01, are selected as those contributing to the specified singular value vectors. Nevertheless, because such a tensor can be obtained only when drug treatment is performed on patients, this strategy is useless; if we can test drug efficiency directly on patients, then there is no need for *in silico* drug discovery. To overcome this discrepancy, I replace 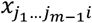 with a product, 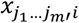· 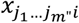, where 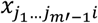 is gene expression for the drug treatment of cell lines/model animals, while 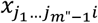 is gene expression for the patients (*m* − 1 = *m*′ + *m*″). Because these two can be obtained independently, we can test any kind of combinations of drug treatments and diseases even after all measurements were performed.

The process performed in the present study following the above procedures is illustrated in Figure 1.

**Figure 1.**
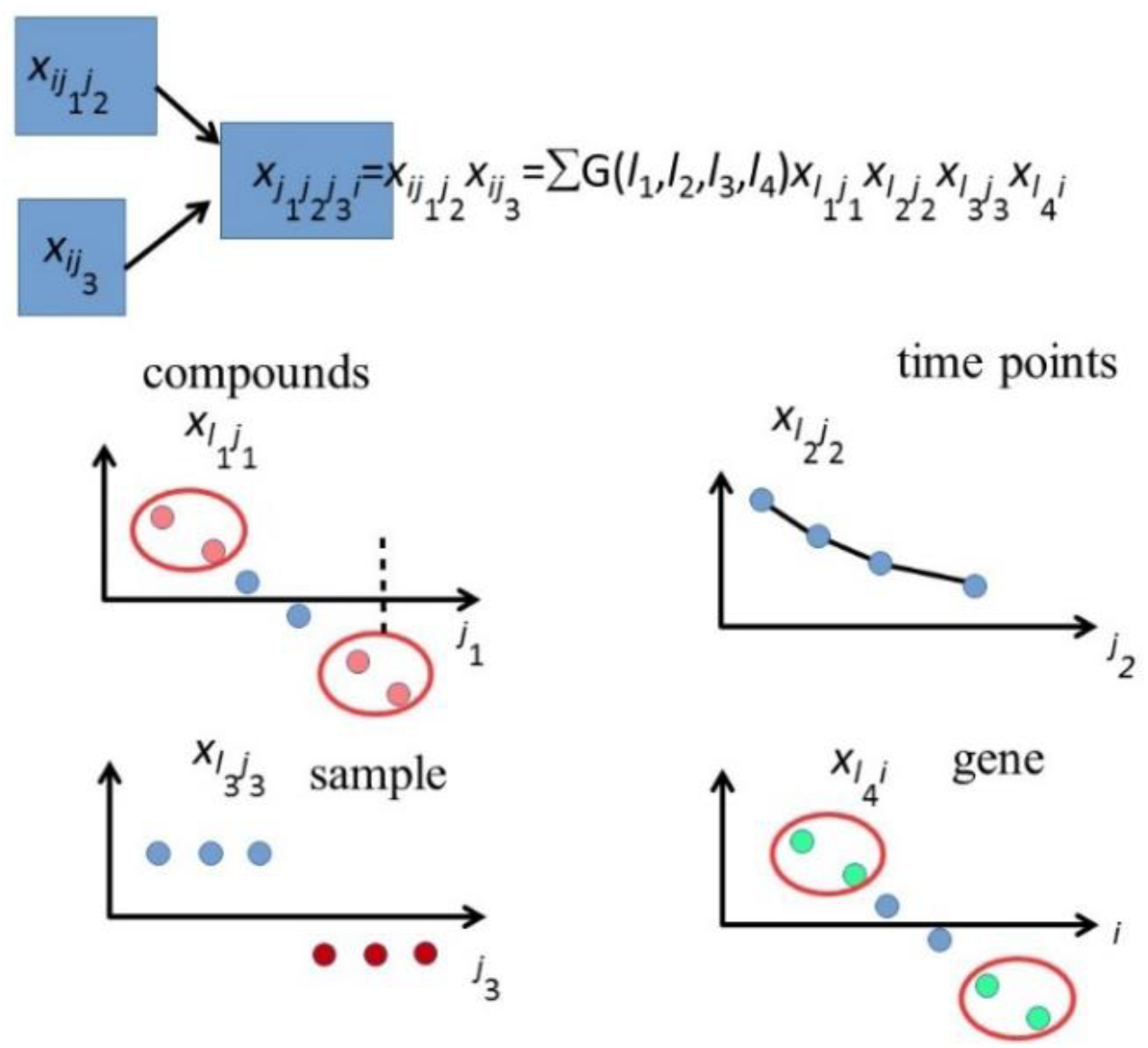
Schematic that illustrates gene-drug pairs identification processes performed in the present study. Gene expression from DrugMatrix, 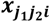, and heart failure, 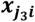, are multiplied and tensor, 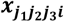, is generated. It is decomposed into core matrix, *G*, and compound singular value matrix 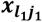, time point singular value matrix, 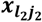, human sample singular value matrix, 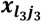, and gene singular value matrix, 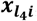, are generated. After the temporal dependent 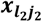 and disease dependent 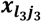 are identified, associated genes and compounds 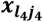 and 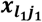 are selected. Based upon them, genes and compounds as outliers are selected by assuming a *χ*^2^ distribution.

### 2.3 Gene Expression Profiles

Gene expression profiles for drug treatments of rats were retrieved from Drug Matrix under the gene expression omnibus (GEO) ID GSE59905, while heart failure human gene expression was taken from GEO ID 57345. For both datasets, expression files of genes, GSE57345-GPL11532_series_matrix.txt.gz, GSE59905GPL5426_series_matrix.txt.gz, and GSE59905-GPL5425_series_matrix.txt.gz were directly downloaded from the series matrix.

### 2.4 Various Servers for Enrichment Analysis

To Enrichr [33] and TargetMine [34], 274 gene symbols were uploaded. For TargetMine, human was assumed as an organism under study, and the BH criterion was used for *P*-value correction.

### 2.5 Statistical Analysis

All the statistical analyses were performed within the R software. HOSVD was carried out using the `hosvd` function in the rTensor package.

## 3 Results

### 3.1 TD-based Unsupervised FE Was Applied to a Combined Tensor

From gene expression profiles of the rat left ventricle (LV) treated with 218 drugs, we selected four time points (1/4, 1, 3, and 5 days after treatment). Although these do not directly represent drug dose dependence, time course observations can be replaced with dose dependence, because drug dose density is expected to monotonically decrease with time. On the other hand, human heart gene expression profiles are composed of 82 idiopathic dilated cardiomyopathy patients, 95 ischemic patients, and 136 healthy controls, respectively. Among them, 3937 genes sharing gene symbols between human and rat were considered. Then, the generated tensor is

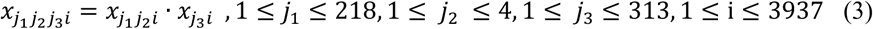

which represents the products of gene expression of the *i*th gene of LV treated with *j*_1_ compound at the *j*_2_ th time point after the drug treatment and gene expression of the *j*_3_ th human heart, respectively. HOSVD was applied to 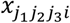 and core tensor *G*(*l*_1_*l*_2_*l*_3_*l*_4_), 1 ≤ *l*_1_ ≤ 218, 1 ≤ *l*_2_ ≤ 4, 1 ≤ *l*_3_ ≤ 313, 1 ≤ *l*_4_ ≤ 3937, compound singular value matrix 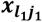, time point singular value matrix, 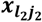, human sample singular value matrix, 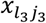, and gene singular value matrix, 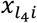, were obtained. Prior to selection of genes and compounds, we need to know which time points singular value vector represents time dependence and which human sample singular value vector represents the distinction between patients and healthy controls (Fig. 2). As for time point singular value vectors, I decided to use the 2^nd^ time point singular value vector, 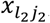, *l*_2_ = 2, because it has the strongest correlation with days. It also represents reasonable time development. After drug treatment, gene expression gradually increases because it takes a while for a drug treatment to have an effect. Then, after it has a peak on day 1, a monotonic decrease follows. On the other hand, for human sample singular value vectors, the 2^nd^ and 3^rd^ ones were selected because they have a clear distinction between patients and healthy controls.

**Figure 2 Left:**
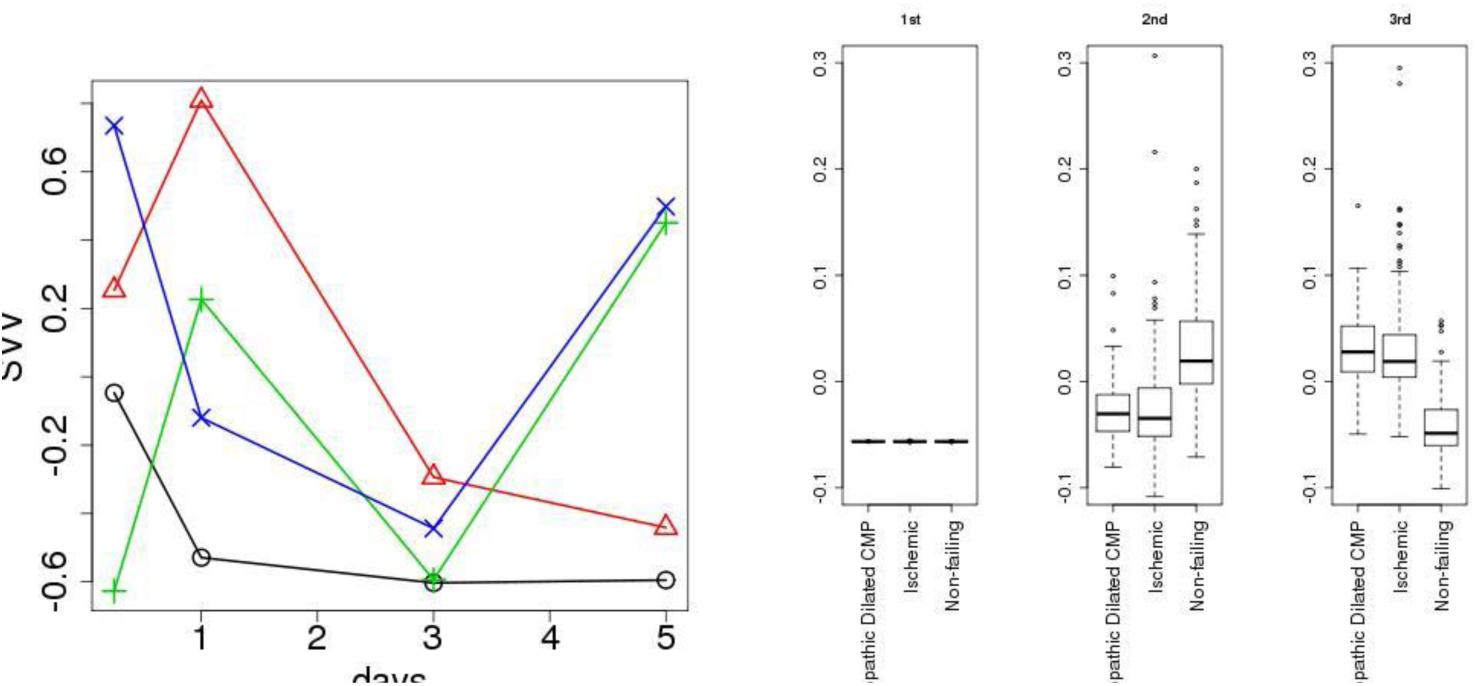
Time points’ singular value vectors. Black circle: 1^st^, red triangle: 2^nd^, green cross: 3^rd^, and blue cross: 4^th^ singular value vectors, respectively. Pearson’s correlation coefficients toward days are -0.72, -0.82, 0.51, and -0.09, respectively. **Right**: A box plot of human sample singular value vectors. From left to right, the 1^st^, 2^nd^, and 3^rd^ singular value vectors are shown.

Next, I tried to identify gene singular value vectors and compound singular value vectors associated with core tensor *G*(*l*_1_*l*_2_*l*_3_*l*_4_), *l*_2_ = 2, 2 ≤ *l*_3_ ≤ 3 that have larger absolute values (Table 1). One can see that the 2^nd^ singular value vector of compounds is always associated with top 20 core tensors. The selection of gene singular value vectors is not so trivial. First of all, generally low-ranked gene singular value vectors are listed. This means that gene expression associated with disease progression is not a majority. This is a common situation because the disease usually affects only a limited number of genes. Then, tentatively, I decided to select top 10 gene singular value vectors, 21^st^, 25^th^, 27^th^, 28^th^, 33^rd^, 36^th^, 37^th^, 38^th^, 41^st^, and 42^nd^ singular value vectors of genes. Using these singular value vectors, *P* values were attributed to genes and compounds. The attributed *P* values were adjusted by the BH criterion. Then, 281 probes and 0 compounds associated with adjusted *P* values less than 0.01 were selected. Because no compounds pass our criteria, I sought another way to select compounds. Fig. 3 shows the histogram of the 2^nd^ singular value vectors of compounds. There are obviously some outliers. Then, tentatively, I selected 43 compounds having the absolute 2^nd^ singular value vector components larger than 0.1.

**Table 1.**
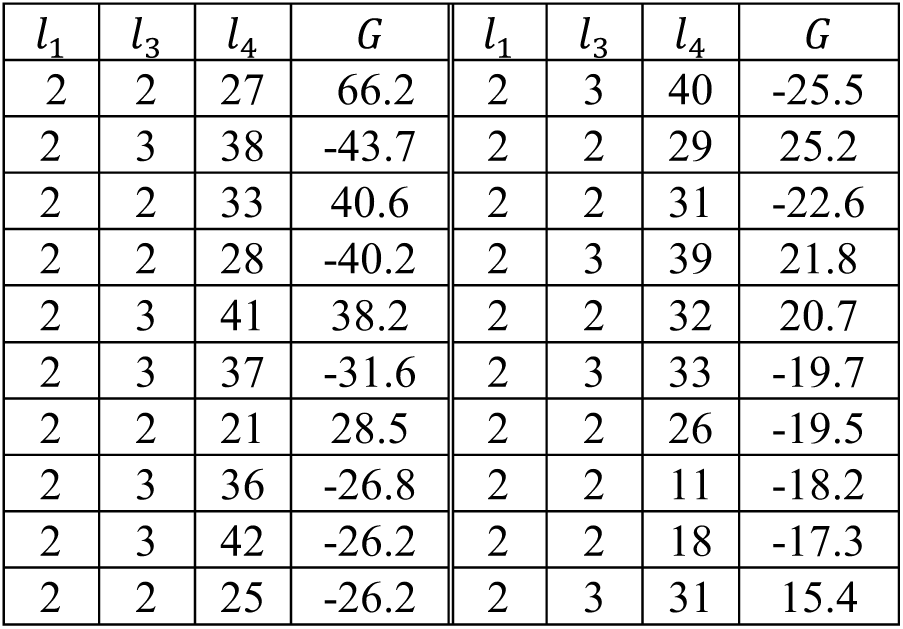
*G*(*l*_1_*l*_2_*l*_3_*l*_4_), *l*_2_ = 2, 2 ≤ *l*_3_ ≤ 3, in the order of larger absolute values of *G*

| $l_1$ | $l_3$ | $l_4$ | $G$ | $l_1$ | $l_3$ | $l_4$ | $G$ |
| --- | --- | --- | --- | --- | --- | --- | --- |
| 2 | 2 | 27 | 66.2 | 2 | 3 | 40 | -25.5 |
| 2 | 3 | 38 | -43.7 | 2 | 2 | 29 | 25.2 |
| 2 | 2 | 33 | 40.6 | 2 | 2 | 31 | -22.6 |
| 2 | 2 | 28 | -40.2 | 2 | 3 | 39 | 21.8 |
| 2 | 3 | 41 | 38.2 | 2 | 2 | 32 | 20.7 |
| 2 | 3 | 37 | -31.6 | 2 | 3 | 33 | -19.7 |
| 2 | 2 | 21 | 28.5 | 2 | 2 | 26 | -19.5 |
| 2 | 3 | 36 | -26.8 | 2 | 2 | 11 | -18.2 |
| 2 | 3 | 42 | -26.2 | 2 | 2 | 18 | -17.3 |
| 2 | 2 | 25 | -26.2 | 2 | 3 | 31 | 15.4 |

**Figure 3.**
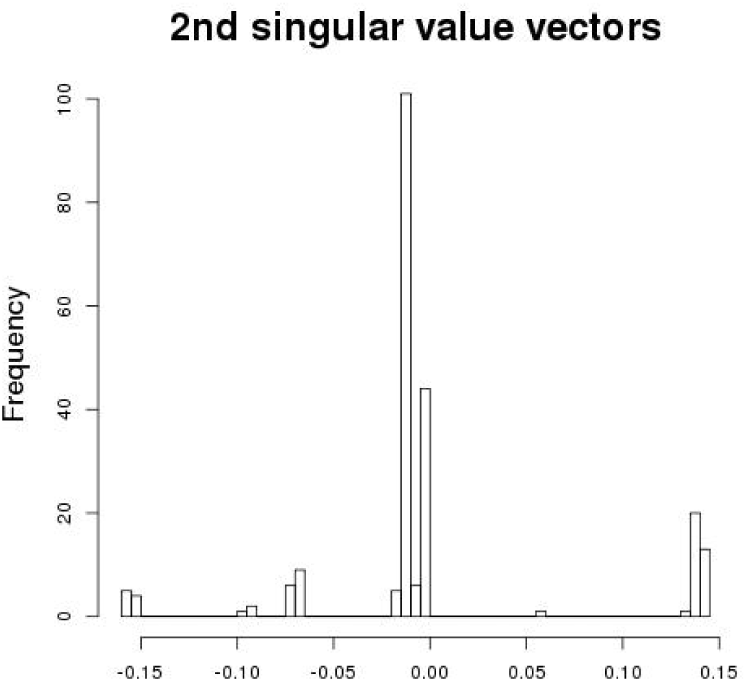
A histogram of 2^nd^ singular value vectors of compounds.

### 3.2 Biological Evaluation of the Selected Compounds and Genes

To see if we can successfully identify biologically relevant compounds and genes, we evaluated these selected genes and compounds. At first, a literature search was performed on the 43 drugs. Then, some heart failure-related studies were identified for most of the 43 drugs (Table 2). This means that biologically relevant drugs were likely to be identified successfully. As for the genes identified, 274 genes associated with the identified 281 probes are shown in Table 3.

**Table 2.** Literature search performed on 43 drugs identified by TD-based unsupervised FE. Numbers are Pubmed IDs (https://www.ncbi.nlm.nih.gov/pubmed/) that report the relation to heart failure.

|  |  |  |  |  |
| --- | --- | --- | --- | --- |
| Amitriptyline<br>[27994924] | Atropine<br>[24279866] | Baclofen<br>[27682809] | Bezafibrate<br>[26957517] | Caffeine<br>[25944789] |
| Calcitriol<br>[27209698] | Chlorambucil<br>[8164221] | Cimetidine<br>[18656805] | Citalopram<br>[25326372] | Clemastine<br>[16288909] |
| Clonazepam<br>[15699937] | Cyclophosphamide<br>[24467219] | D-Tubocurarine<br>Chloride<br>[14839919] | Dexamethasone<br>[25923220] | Dexchlor-<br>pheniramine<br>[---] |
| Digitoxin<br>[27082032] | Diphenhydramine<br>[22158278] | Doxazosin<br>[26515144] | Ebastine<br>[21410688] | Fenofibrate<br>[26497978] |
| Fluphenazine<br>[23395964] | Gabapentin<br>[19195912] | Ifosfamide<br>[14586140] | Iproniazid<br>[13688979] | Lacidipine<br>[23911888] |
| Loratadine<br>[21880544] | Nevirapine<br>[15526045] | Nimodipine<br>[17191657] | Nitrendipine<br>[22750214] | Ofloxacin<br>[21559378] |
| Oxymetazoline<br>[22855901] | Paroxetine<br>[26216863] | Phenacemide<br>[---] | Phenytoin<br>[24172819] | Rosiglitazone<br>[21666037] |
| Sparteine [4408029] | Stavudine [---] | Valsartan<br>[26992459] | Vecuronium<br>Bromide [---] | Venlafaxine<br>[23301719] |
| Vinblastine<br>[25537132] | Vincristine [---] | Zidovudine<br>[25838291] |  |  |

**Table 3.**
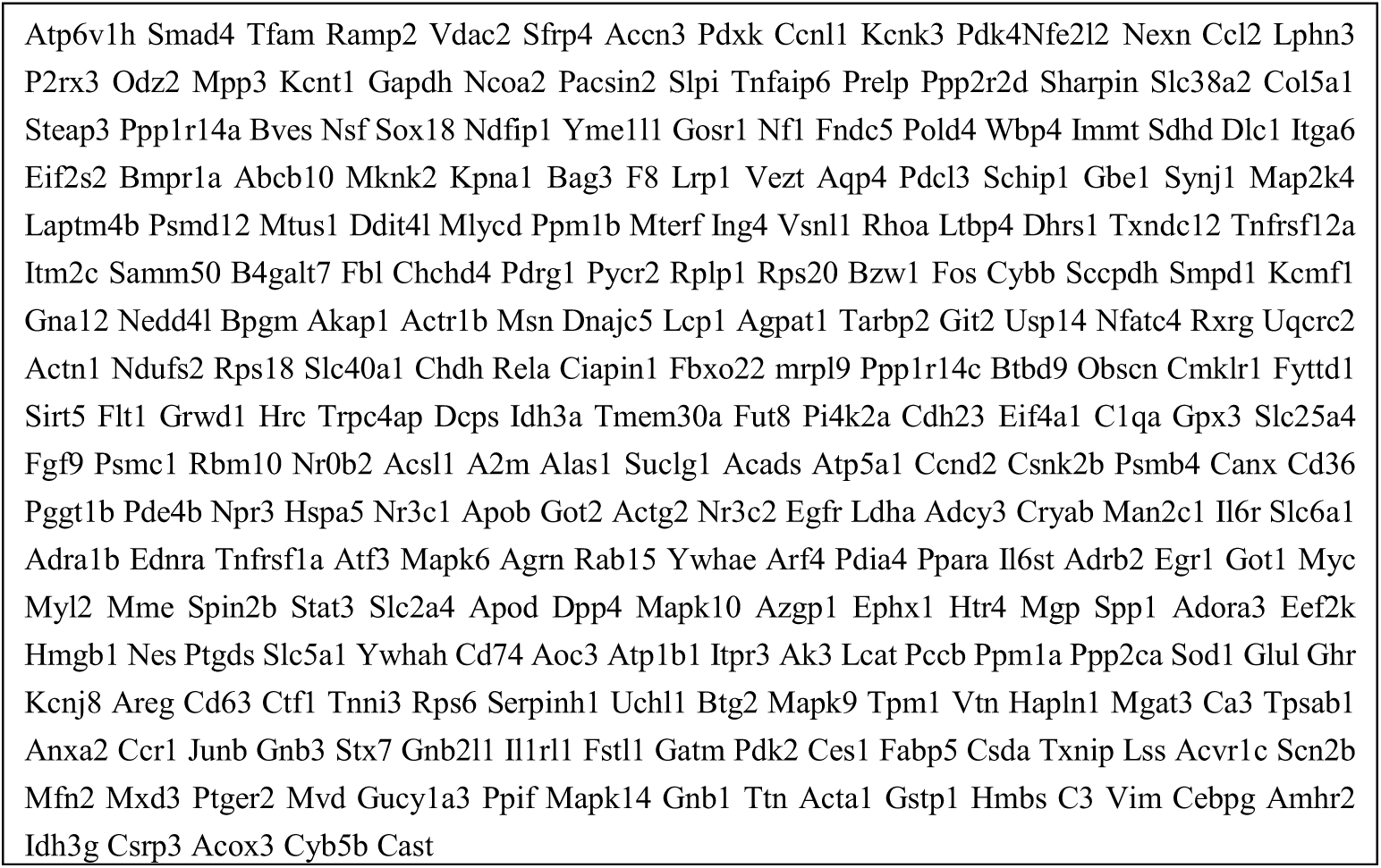
The 274 genes associated with 281 probes identified by TD-based unsupervised FE.

To evaluate biological reliability of these 274 genes, they were uploaded to various enrichment servers. When they were uploaded to TargetMine, top five tissue enrichment results were related to the heart (Table 4). Top four significant disease enrichment results represent heart failure (Table 5). When they were uploaded to Enrichr, top three OMIM disease enrichment results were related to heart failure (Table 6). Two out of top three MGI Mammalian Phenotype Level 3 enrichment results were also related to heart failure (Table 7). Thus, our identification of genes was also successful.

**Table 4.** Top five significant tissue enrichment results of TargetMine.

| Tissue | *p*-value | Matches |
| --- | --- | --- |
| left ventricular apex samples | 1.880e-37 | 101 |
| heart atrium | 1.357e-36 | 155 |
| heart | 8.637e-36 | 153 |
| ventricular myocardium | 4.306e-35 | 95 |
| atrial myocardium | 1.716e-34 | 98 |

**Table 5.**
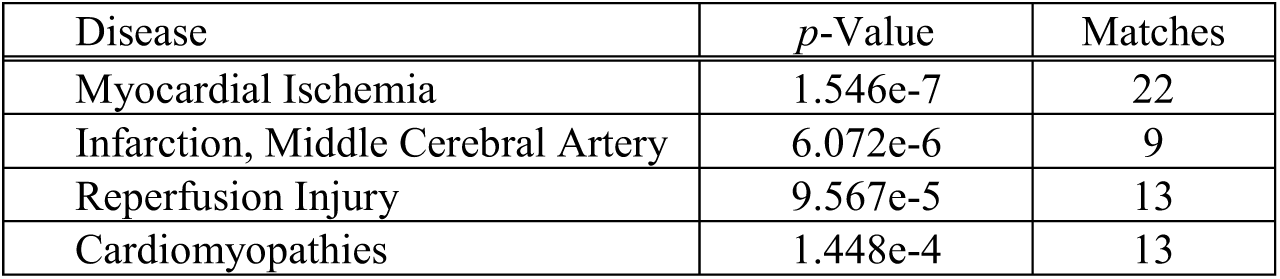
Top four disease enrichment results of TargetMine.

**Table 6.** Top three significant OMIM Disease enrichment results of Enrichr.

| Name | Overlap | *P*-value | Adjusted *p*-value |
| --- | --- | --- | --- |
| cardiomyopathy, hypertrophic | 5/17 | 2.516E-05 | 9.814E-05 |
| cardiomyopathy | 5/42 | 2.615E-04 | 5.099E-03 |
| cardiomyopathy, dilated | 4/33 | 1.031E-03 | 1.341E-02 |

**Table 7.**
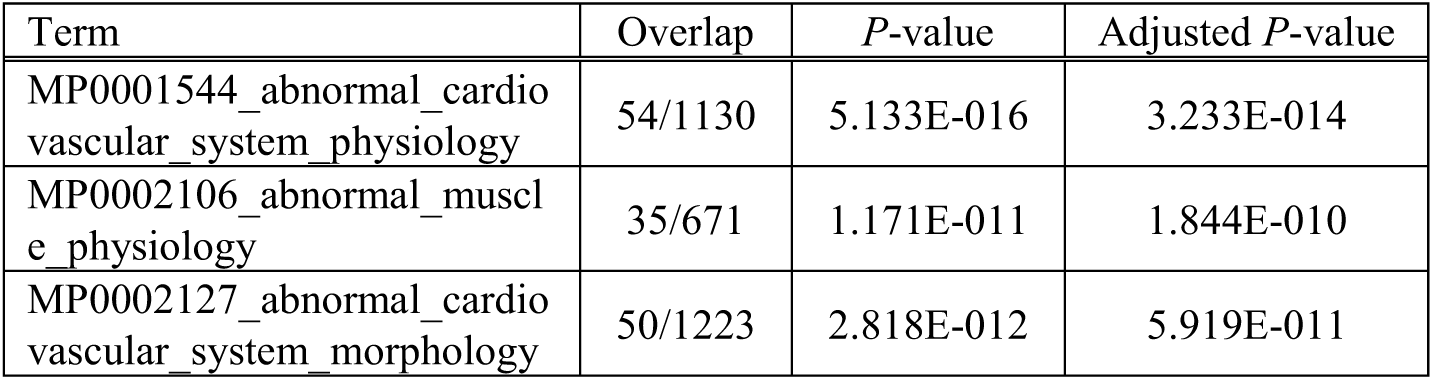
Top three significant MGI Mammalian Phenotype Level 3 enrichment results of Enrichr.

| Term | Overlap | *P*-value | Adjusted *P*-value |
| --- | --- | --- | --- |
| MP0001544_abnormal_cardiovascular_system_physiology | 54/1130 | 5.133E-016 | 3.233E-014 |
| MP0002106_abnormal_muscle_physiology | 35/671 | 1.171E-011 | 1.844E-010 |
| MP0002127_abnormal_cardiovascular_system_morphology | 50/1223 | 2.818E-012 | 5.919E-011 |

Enrichr also outputted many epigenetic feature enrichment results. Top most significant ENCODE TF-ChiP-seq 2015 is POLR2A_heart_mm9; POLR2A is a transcription factor (TF) reported to be a stable reference gene for gene expression alteration in gene expression studies on rodent and human heart failure [35]. This finding suggested that *POLR2A* is constantly expressive in heart failure, which is coincident with our analysis. Top most significant TF-LOF Expression result from GEO is yy1_227711985_skeletal_muscle_lof_mouse_gpl8321_gse39009_up. YY1 is a TF reported to play critical roles in cardiac morphogenesis [36]. The top most significant ENCODE Histone Modifications 2015 result is H3K36me3_myocyte_mm9; H3K36me3 was reported to play a crucial role in cardiomyocyte differentiation [37]. These TFs as well as histone modifications identified by our strategy can be possible drug targets.

## 4 Conclusions

In this paper, I introduced a new strategy that integrates disease (heart failure) gene expression profiles with drug treatment-related tissue gene expression profiles. The identified genes as well as compounds have been widely reported to be related to heart failure. Thus, this strategy turned out to be useful for *in silico* drug discovery.

## Footnotes

1 https://ntp.niehs.nih.gov/drugmatrix/index.html

## References

1. Favia, A.D.: Theoretical and computational approaches to ligand-based drug discovery. Front. Biosci. (Landmark Ed. 16, 1276–90 (2011).

2. Lionta, E., Spyrou, G., Vassilatis, D., Cournia, Z.: Structure-Based Virtual Screening for Drug Discovery: Principles, Applications and Recent Advances. Curr. Top. Med. Chem. 14, 1923–1938 (2014).

3. Liu, C., Su, J., Yang, F., Wei, K., Ma, J., Zhou, X.: Compound signature detection on LINCS L1000 big data. Mol. BioSyst. 11, 714–722 (2015).

4. Hizukuri, Y., Sawada, R., Yamanishi, Y.: Predicting target proteins for drug candidate compounds based on drug-induced gene expression data in a chemical structure-independent manner. BMC Med. Genomics. 8, 82 (2015).

5. Wang, K., Sun, J., Zhou, S., Wan, C., Qin, S., Li, C., He, L., Yang, L.: Prediction of Drug-Target Interactions for Drug Repositioning Only Based on Genomic Expression Similarity. PLoS Comput. Biol. 9, e1003315 (2013).

6. Iwata, M., Sawada, R., Iwata, H., Kotera, M., Yamanishi, Y.: Elucidating the modes of action for bioactive compounds in a cell-specific manner by large-scale chemically-induced transcriptomics. Sci. Rep. 7, 40164 (2017).

7. Lee, H., Kang, S., Kim, W., Fedorov, O., Filippakopoulos, P., Hunt, J.: Drug Repositioning for Cancer Therapy Based on Large-Scale Drug-Induced Transcriptional Signatures. PLoS One. 11, e0150460 (2016).

8. Cheng, J., Yang, L., Kumar, V., Agarwal, P.: Systematic evaluation of connectivity map for disease indications. Genome Med. 6, 95 (2014).

9. Sirota, M., Dudley, J.T., Kim, J., Chiang, A.P., Morgan, A.A., Sweet-Cordero, A., Sage, J., Butte, A.J.: Discovery and Preclinical Validation of Drug Indications Using Compendia of Public Gene Expression Data. Sci. Transl. Med. 3, (2011).

10. Iorio, F., Bosotti, R., Scacheri, E., Belcastro, V., Mithbaokar, P., Ferriero, R., Murino, L., Tagliaferri, R., Brunetti-Pierri, N., Isacchi, A., di Bernardo, D.: Discovery of drug mode of action and drug repositioning from transcriptional responses. Proc. Natl. Acad. Sci. U. S. A. 107, 14621–6 (2010).

11. Kinoshita, R., Iwadate, M., Umeyama, H., Taguchi, Y.H.: Genes associated with genotype-specific DNA methylation in squamous cell carcinoma as candidate drug targets. BMC Syst Biol. 8 Suppl 1, S4 (2014).

12. Taguchi, Y., Iwadate, M., Umeyama, H., Murakami, Y., Okamoto, A.: Heuristic principal component analysis-aased unsupervised feature extraction and its application to bioinformatics. In: Wang, B., Li, R., and Perrizo, W. (eds.) Big Data Analytics in Bioinformatics and Healthcare. pp. 138–162 (2015).

13. Murakami, Y., Kubo, S., Tamori, A., Itami, S., Kawamura, E., Iwaisako, K., Ikeda, K., Kawada, N., Ochiya, T., Taguchi, Y.H.: Comprehensive analysis of transcriptome and metabolome analysis in Intrahepatic Cholangiocarcinoma and Hepatocellular Carcinoma. Sci Rep. 5, 16294 (2015).

14. Taguchi, Y.-H., Iwadate, M., Umeyama, H.: Heuristic principal component analysis-based unsupervised feature extraction and its application to gene expression analysis of amyotrophic lateral sclerosis data sets. 2015 IEEE Conf. Comput. Intell. Bioinforma. Comput. Biol. (2015).

15. Umeyama, H., Iwadate, M., Taguchi, Y.: TINAGL1 and B3GALNT1 are potential therapy target genes to suppress metastasis in non-small cell lung cancer. BMC Genomics. 15, S2 (2014).

16. Murakami, Y., Kubo, S., Tamori, A., Itami, S., Kawamura, E., Iwaisako, K., Ikeda, K., Kawada, N., Ochiya, T., Taguchi, Y.-H.: Comprehensive analysis of transcriptome and metabolome analysis in Intrahepatic Cholangiocarcinoma and Hepatocellular Carcinoma. Sci. Rep. 5, 16294 (2015).

17. Taguchi, Y., Murakami, Y.: Principal component analysis based feature extraction approach to identify circulating microRNA biomarkers. PLoS One. (2013).

18. Taguchi, Y.-H., Murakami, Y.: Universal disease biomarker: can a fixed set of blood microRNAs diagnose multiple diseases? BMC Res. Notes. 7, 581 (2014).

19. Murakami, Y., Tanahashi, T., Okada, R., Toyoda, H., Kumada, T., Enomoto, M., Tamori, A., Kawada, N., Taguchi, Y.H., Azuma, T.: Comparison of hepatocellular carcinoma miRNA expression profiling as evaluated by next generation sequencing and microarray. PLoS One. 9, (2014).

20. Taguchi, Y. -h., Iwadate, M., Umeyama, H.: Principal component analysis-based unsupervised feature extraction applied to in silico drug discovery for posttraumatic stress disorder-mediated heart disease. BMC Bioinformatics. 16, 139 (2015).

21. Y-h. Taguchi: Identification of More Feasible MicroRNA–mRNA Interactions within Multiple Cancers Using Principal Component Analysis Based Unsupervised Feature Extraction. Int. J. Mol. Sci. 17, E696 (2016).

22. Taguchi, Y.-H., Iwadate, M., Umeyama, H.: Heuristic principal component analysis-based unsupervised feature extraction and its application to gene expression analysis of amyotrophic lateral sclerosis data sets. In: Computational Intelligence in Bioinformatics and Computational Biology (CIBCB), 2015 IEEE Conference on. pp. 1–10 (2015).

23. Taguchi, Y.-H.: Principal component analysis based unsupervised feature extraction applied to publicly available gene expression profiles provides new insights into the mechanisms of action of histone deacetylase inhibitors. NEPIG. (2016).

24. Taguchi, Y.-H., Iwadate, M., Umeyama, H.: SFRP1 is a possible candidate for epigenetic therapy in non-small cell lung cancer. BMC Med. Genomics. 9, (2016).

25. Taguchi, Y.-H.: microRNA-mRNA interaction identification in Wilms tumor using principalcomponent analysis based unsupervised feature extraction. In: 2016 IEEE 16th International Conference on Bioinformatics and Bioengineering (BIBE). pp. 71–78 (2016).

26. Taguchi, Y.-H.: Principal component analysis based unsupervised feature extraction applied to budding yeast temporally periodic gene expression. BioData Min. 9, 22 (2016).

27. Murakami, Y., Toyoda, H., Tanahashi, T., others: Comprehensive miRNA expression analysis in peripheral blood can diagnose liver disease. PLoS One. 7, e48366 (2012).

28. Ishida, S., Umeyama, H., Iwadate, M., Taguchi, Y.H.: Bioinformatic Screening of Autoimmune Disease Genes and Protein Structure Prediction with FAMS for Drug Discovery. Protein Pept. Lett. 21, 828–839 (2014).

29. Taguchi, Y.: Principal Components Analysis Based Unsupervised Feature Extraction Applied to Gene Expression Analysis of Blood from Dengue Haemorrhagic Fever Patients. Sci. Rep. 7, 44016 (2017).

30. De Lathauwer, L., De Moor, B., Vandewalle, J.: A Multilinear Singular Value Decomposition. SIAM J. Matrix Anal. Appl. 21, 1253–1278 (2000).

31. Duan, Q., Reid, S.P., Clark, N.R., Wang, Z., Fernandez, N.F., Rouillard, A.D., Readhead, B., Tritsch, S.R., Hodos, R., Hafner, M., Niepel, M., Sorger, P.K., Dudley, J.T., Bavari, S., Panchal, R.G., Ma’ayan, A.: L1000CDS2: LINCS L1000 characteristic direction signatures search engine. npj Syst. Biol. Appl. 2, 16015 (2016).

32. Benjamini, Y., Hochberg, Y.: Controlling the false discovery rate: a practical and powerful approach to multiple testing. J. R. Stat. Soc. B57, 289–300 (1995).

33. Kuleshov, M. V., Jones, M.R., Rouillard, A.D., Fernandez, N.F., Duan, Q., Wang, Z., Koplev, S., Jenkins, S.L., Jagodnik, K.M., Lachmann, A., McDermott, M.G., Monteiro, C.D., Gundersen, G.W., Ma’ayan, A.: Enrichr: a comprehensive gene set enrichment analysis web server 2016 update. Nucleic Acids Res. 44, W90–W97 (2016).

34. Chen, Y.-A., Tripathi, L.P., Mizuguchi, K.: TargetMine, an Integrated Data Warehouse for Candidate Gene Prioritisation and Target Discovery. PLoS One. 6, e17844 (2011).

35. Brattelid, T., Winer, L.H., Levy, F.O., Liestøl, K., Sejersted, O.M., Andersson, K.B.: Reference gene alternatives to Gapdh in rodent and human heart failure gene expression studies. BMC Mol. Biol. 11, 22 (2010).

36. Beketaev, I., Zhang, Y., Kim, E.Y., Yu, W., Qian, L., Wang, J.: Critical role of YY1 in cardiac morphogenesis. Dev. Dyn. 244, 669–680 (2015).

37. Cattaneo, P., Kunderfranco, P., Greco, C., Guffanti, A., Stirparo, G.G., Rusconi, F., Rizzi, R., Di Pasquale, E., Locatelli, S.L., Latronico, M.V.G., Bearzi, C., Papait, R., Condorelli, G.: DOT1L-mediated H3K79me2 modification critically regulates gene expression during cardiomyocyte differentiation. Cell Death Differ. 23, 555–64 (2016).

